# Computational design of highly signaling active membrane receptors through *de novo* solvent-mediated allosteric networks

**DOI:** 10.1101/2021.09.29.462228

**Authors:** K-Y. Chen, J.K. Lai, J. Wang, A.M. Russell, K. Conners, M.E. Rutter, B. Condon, F. Tung, L. Kodandapani, B. Chau, X. Zhao, J. Benach, K. Baker, E.J. Hembre, P. Barth

## Abstract

Protein catalysis and allostery require the atomic-level orchestration and motion of residues, ligand, solvent and protein effector molecules, but the ability to design protein activity through precise protein-solvent cooperative interactions has not been demonstrated. Here, we report the design of a dozen novel membrane receptors catalyzing G-protein nucleotide exchange through diverse *de novo* engineered allosteric pathways mediated by cooperative networks of intra-protein, protein-ligand and solvent molecule interactions. Consistent with the predictions, designed protein activities strongly correlated with the level of solvent-mediated interaction network plasticity at flexible transmembrane helical interfaces. Several designs displayed considerably enhanced thermostability and activity compared to related natural receptors. The most stable and active variant crystallized in an unforeseen signaling active conformation, in excellent agreement with the design models. The allosteric network topologies of the best designs bear limited similarity to those of natural receptors and reveal a space of allosteric interactions larger than previously inferred from natural proteins. The approach should prove useful for engineering proteins with novel complex protein catalytic and signaling activities.

## Introduction

The remarkable efficiency and selectivity observed in natural protein catalysts, binders and signaling receptors result from a delicate balance between stabilizing features that define the molecular structure and destabilizing functional motifs. For example, exposed hydrophobic surfaces in soluble proteins encode binding affinity and specificity but can trigger protein aggregation. Enzymes often bear energetically unfavorable networks of buried polar residues in their active sites that are critical for high-precision substrate binding and catalytic activity. Membrane proteins perform sophisticated signaling and transport functions by rapidly switching between conformations and enabling long-range allosteric communication between both sides of the membrane^1^. Wet transmembrane helical (TMH) interfaces where networks of solvent molecules bridge destabilizing buried polar residues in membrane protein cores, facilitate TMH movements by preventing the breaking of hydrogen-bond networks^2,3^. On the other hand, buried ion molecules decrease conformational flexibility by locking receptors in selective conformations and functional states through strong electrostatic interactions^4^. Similarly, intricate networks of buried and conserved solvent-mediated polar interactions in protein kinases define allosteric pathways that directly control substrate binding cooperativity^5^. Hence, cooperative protein-solvent interaction networks have emerged as key universal principles of protein function.

Computational methods that incorporate typical features of natural protein catalysts and binders in protein designs have been developed in recent years. A high level of structural specificity and stability was achieved in helical bundles through de novo designed geometrically optimized hydrogen-bond networks^6,7^. Key structural features that constitute the hallmark of natural enzymes, including hydrophobic cavities, kinked helices and curved sheets, can now be combined to create novel protein scaffolds^8,9^. These approaches enable an unprecedented level of control over biomolecular shape and interactions. However, leveraging the delicate balance between stabilizing and destabilizing functional features to design new protein activities remains a major challenge.

Here, we report a computational approach that enables the efficient and accurate design of proteins with novel stability and signaling functions through *de novo* engineered cooperative intra-protein, protein-ligand and protein-solvent molecule interaction networks. Starting from the scaffold of a classical G-protein coupled receptor (GPCR), we applied the method to design membrane receptors that achieved a wide range of signaling activity and thermostability. In close agreement with the design models, a highly signaling active designed receptor crystallized in an atypical GPCR active-like conformation despite the absence of a bound G-protein. Our results reveal that signaling activity and thermostability can be engineered concurrently through a careful modulation of reprogrammable ligand and solvent-mediated allosteric interaction networks. Our approach should prove useful for engineering both protein catalysts and membrane receptor biosensors with new signal transduction functions, and accelerating membrane receptor structure determination.

## Results

### Design principles of solvent-mediated allosteric networks

Protein catalysis and allostery often rely on the concerted motions of flexible regions that switch between distinct conformations around a stable and rigid scaffold. Interactions between flexible and rigid structures are defined as “static-switchable” in our study. By bridging buried polar residues and creating extensive networks of dynamic interactions, water molecules can facilitate such movements in transmembrane receptors. By contrast, ion molecules tend to stabilize specific conformations through strong electrostatic interactions at the expense of protein flexibility. Following these observations, we reasoned that signaling activity could be programmed into membrane receptor scaffolds by designing solvent-mediated interaction networks connecting switchable TMH-TMH interfaces that are critical for activation.

To facilitate the design of such properties, we developed a computational method that builds *de novo* protein interiors with customized solvent mediated interaction networks. The method first builds receptor conformations in specific (e.g. inactive and active) functional states by homology to native protein structures with related functions. Rigid and flexible TMH regions are identified from these models, and the interfaces in between are then designed using the software SPaDES developed to model and design protein structures and interactions with explicit solvent^10^. In our calculations, SPaDES was implemented into a multi-state design framework to search for combinations of aminoacids and associated solvent molecules that form highly coordinated networks of solvent-mediated interactions between the TMHs in different states (**Fig. 1a,b**). Designs were generated by modulating the following three features: 1) the level of water-mediated hydrogen bond connectivity between static and switchable TMHs that controls the plasticity of the interface; 2) the strength of protein-ion interactions locking and stabilizing the protein into a specific conformation; 3) the overall conformational stability of the active versus that of the inactive state. For example, high levels of protein activity and active state stability can be achieved concurrently by designing dynamic static-switchable interfaces in the active state (**Fig. 1c**) while preventing ion locks in the inactive state (**Fig. 1d**).

**Figure 1.**
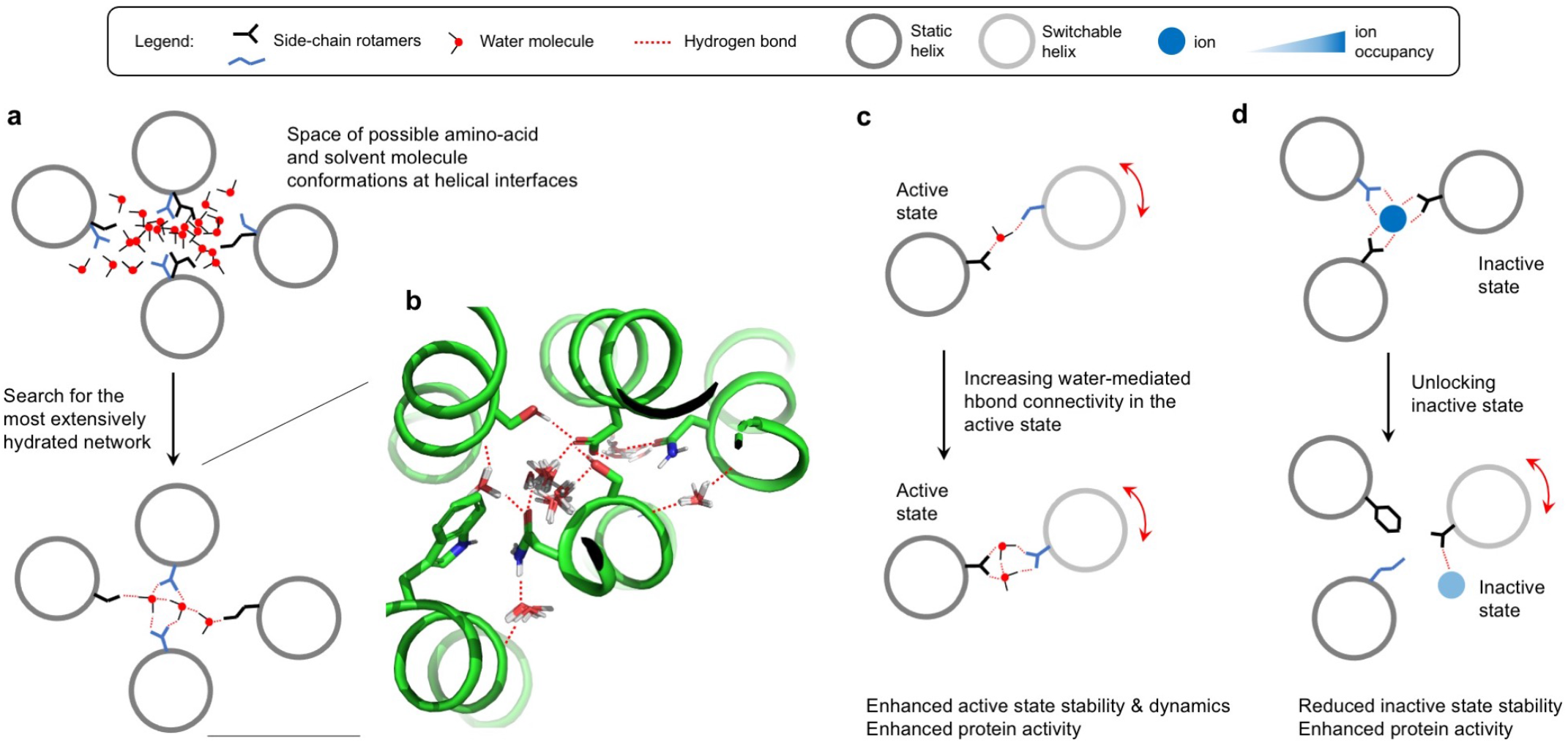
Rational design of signaling activity through *de novo* solvent-mediated interaction networks. **a.** Schematic representation of the design of water-mediated hydrogen bond network bridging 4 helices using SPaDES. *De novo* amino-acid sequences, conformations and water positions are searched concurrently for optimal water-mediated hydrogen bond connectivity at the helical interface. **b.** Snapshot of a low-energy water-mediated hydrogen bond network ensemble in the core of a GPCR structure. **c.** Rationale for enhancing protein activity through increased water-mediated dynamic contacts in the active state. **d.** Rationale for enhancing protein activity through weakening ion-mediated locks in the inactive state.

As a proof of concept, we applied the approach to design signaling activity into an adenosine-sensing GPCR. GPCRs are composed of a topologically conserved 7 transmembrane helical scaffold (7TM) connected by loop regions on either side of the membrane that recognize extracellular ligands and intracellular signaling proteins (e.g. G-proteins, arrestins). The 7TM region transmits extracellular ligand-induced signals through specific concerted motions of TMH6 and TMH7 that trigger the opening of a G-protein binding site on the intracellular side of the receptor (**Fig. 2a**).

**Figure 2.**
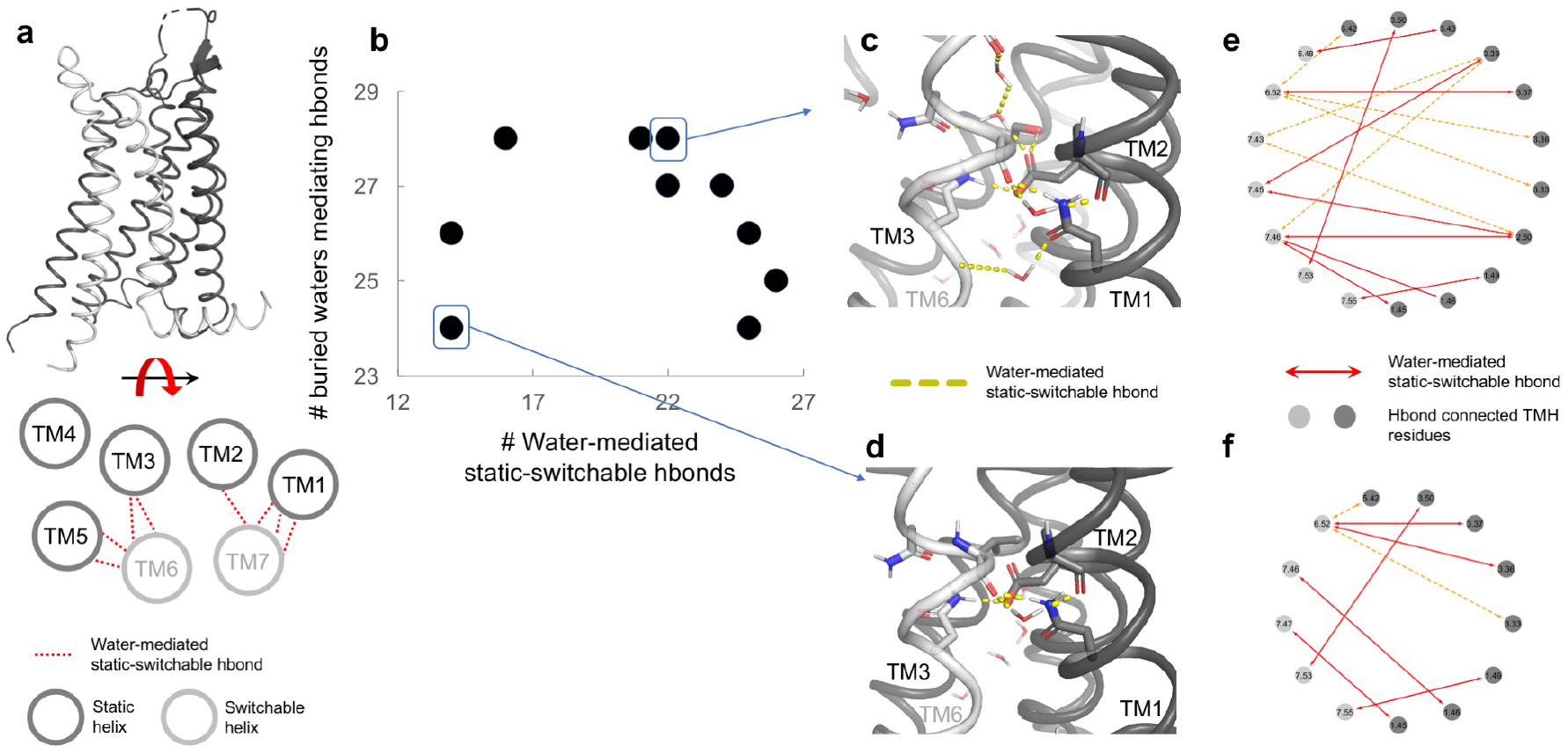
Computational designed GPCRs with a wide range of water-mediated interaction networks at switchable TMH interfaces. **a.** Backbone representation of a GPCR scaffold highlighting static and switchable TM helices. **b.** Relationship between the total number of buried water molecules mediating polar contacts and those at static-switchable TMH interfaces in 10 designed GPCRs. **c-d.** Hydrogen bond network in the TMH core regions of SPaDES_high2 (**c**) and SPaDES_low1 (**d**) structures. **e-f.** Network representation using Cytoscape of the water-mediated hydrogen bonds at static-switchable TMHs of SPaDES_high2 (**e**) and SPaDES_low1 (**f**). Amino-acids are represented as nodes (black on static TMHs, grey on switchable TMHs) with Ballesteros Weinstein (BW) residue notation. Water-mediated hydrogen bonds are represented as colored arrow edges.

Inactive and active state adenosine-sensing GPCR models were constructed from the antagonist bound A2AR X-ray structures and the active state X-ray structure of B2AR bound to the G-protein Gs, respectively. We then applied SPaDES to *de novo* design sequences at the interface between the static TMHs 1, 2, 3 and 5 and the switchable TMHs 6 and 7 that would readily fold into both inactive and active state conformations. A total of 42 buried positions and associated solvent molecules were designed in the TM region of the receptor, giving a total number of 10^32^ possible amino-acid combinations. GPCRs were selected with distinct levels of active versus inactive state stability and plasticity at static-switchable interfaces.

The designs displayed a large diversity of synthetic polar cores and solvent-mediated networks that often bear limited similarity with TMH interaction patterns found in native GPCRs. Overall, we observed a high number of hydrogen bonds mediated by water molecules (up to 28) in the designed cores, but with no obvious correlations with the number of solvated static-switchable interactions that varied between 14 and 26 (**Fig. 2b**). These results suggest that our approach can concurrently design solvent mediated networks essential for the stability of static TMH-TMH interactions or the plasticity of switchable interfaces. We observed two classes of receptors in our design datasets, that we call SPaDES_low and SPaDES_high. SPaDES_low variants tend to display lower active state stability and number of water-mediated static-switchable interactions than SPaDES_high variants (**Fig. 2b**). Overall, the topology of the TMH interaction networks is very diverse across the entire dataset and reflects the large space of possible solutions to the *de novo* construction of TMH interface contact networks.

Detailed analysis of the designed models revealed that even minor sequence variations can profoundly impact the structure of solvent-mediated TMH-TMH contact networks. For example, despite differing by only two residues, SPaDES_low2 and SPaDES_high4 displayed close to 40% different water-mediated static-switchable contacts (**Fig. 2c-f**). SPaDES_high4 bears a critical polar residue near a highly solvated buried Asp at the TMH2-6 interface, while SPaDES_low2 packs a Val at that same position. The presence of a hydrophobic residue considerably rewires the hydrogen bond network and decreases the overall hydration at that switchable interface. Additionally, a highly flexible Met at position 95 (BW) on TMH3 in SPaDES_high4 shapes a microcavity that enables optimal hydration of the switchable TMH3-6 interface. A Leu at that position in SPaDES_low2 prevents such extended solvent-mediated polar network between the TMHs. Specific designed sequences in SPaDES_high variants tended to also to weaken TMH interactions with ion molecules. For example, carefully designed polar residues near a buried Asp on TMH2 prevented the chelation of a sodium molecule by the Asp in the inactive state. The absence of sodium binding was compensated through less energetically favorable hydrogen bonding to water. These observations are consistent with a higher predicted active state stability for SPaDES_high variants.

We next assessed whether our designed receptors displayed the intended spectrum of signaling activity and active state stability. The adenosine biosensors were expressed in mammalian cells and tested for Gs activation using a reporter for intracellular cAMP production (**Methods**). We first measured constitutive activities in the absence of ligands. Consistent with SPaDES_high variants incorporating more active state stabilizing features, most of these designs displayed higher spontaneous signaling activities than SPaDES_low variants (**Fig. 3a**). To corroborate these results, we also directly assessed receptor conformational stability by measuring receptor activity as a function of incubation time at 37°C. While SPaDES_low variants rapidly unfolded, the half-life of SPaDES_high variants was as high as 45 min (**Fig. 3b,c**). Differences between the two classes of designs were even more pronounced when activities were triggered by adenosine ligand stimulus. Most SPaDES_low designs signaling responses were minimal, while those measured for SPaDES_high variants were strong and similar to naturally evolved adenosine receptors (**Fig. 3d**). These results validated our design concept and suggest that the level of solvent-mediated connectivity at static-switchable TMH interfaces is a key determinant of receptor conformational dynamics and signal transduction propensity.

**Figure 3.**
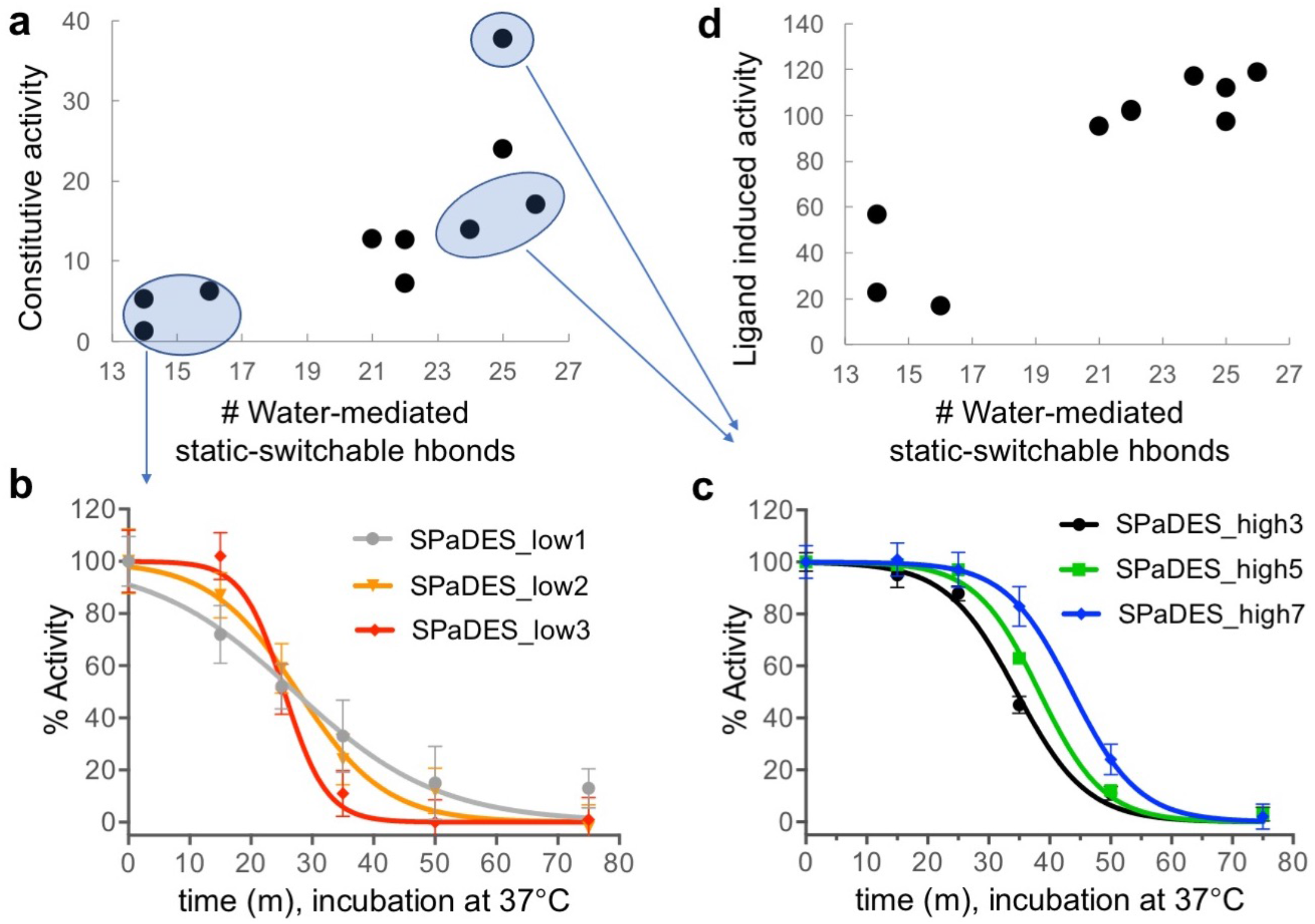
Designed SPaDES stability and function strongly depend on the density of solvent-mediated interactions at switchable TMH interfaces. **a.** Measured constitutive activity of designed SPaDES variants (percentage of adenosine-induced A2AR WT activity) as a function of the number of solvent-mediated interactions at switchable TMH interfaces. **b,c.** Apparent receptor stability measured by the functional half-life of adenosine-bound SPaDES variants incubated at 37°C. **d.** Adenosine-induced activity of designed SPaDES variants (percentage of adenosine-induced A2AR WT activity) as a function of the number of solvent-mediated interactions at switchable TMH interfaces.

To further validate the predicted designed effects on the binding of a sodium ion-molecule in the TMH core and inactive state stability, we performed thermostability assays under distinct buffer conditions using either potassium or sodium as the main monovalent cation. A 4°C increase in thermostability was observed for agonist-bound A2AR WT and designs predicted to not substantially alter Na^+^ binding, following replacement of Na^+^ by K^+^. This is consistent with our calculations and the documented role of Na^+^ as a negative allosteric modulator in native GPCRs^10^, as the increased size of K^+^ prevents it from accessing the same sites as Na^+^. Conversely, the thermostability of SPaDES variants, which incorporated the S91T (i.e. the primary determinant for preventing tight Na^+^ binding to D52), lost dependency to the monovalent cation identity.

We next sought to validate the structure of our designed receptors. We selected our most stabilized variants and screened for potent agonists and fusion proteins at the N terminus of the receptor to achieve maximal expression and apparent agonist-bound stability^11^ (**Methods**). When fused to a circularly permutant T4L variant (T4Lcp) and bound to the CGS21680 agonist, SPaDES_high7 displayed a large increase in thermodynamic stability compared to the native A2AR (close to 20°C), reaching an apparent melting temperature of 72°C. We successfully crystallized the CGS21680-bound T4Lcp-SPaDES_high7 construct and solved its structure at 3.9 Å resolution (**Figure 4a**). While the moderate resolution prevented the unambiguous assignment of electron density to solvent molecules, the ligand agonist and most side-chain conformations could be readily characterized. In broad agreement with the designed model, the agonist-bound SPaDES_high7 adopted an active-like conformation despite the absence of G-protein. The conformations of TM6 and TM7 were similar to those in the active form of opsin bound to a C-terminal peptide of the G-protein Gt^12^, while the presence of the full length G-protein in the ternary active state structure of GPCRs often induces additional distortions in the intracellular tip of TM6^13^ (**Figure 4b**). GPCR activation involves side-chain movements of several highly-conserved residue microswitches, constituting key molecular signatures of receptor signaling^14,15^. Because they are critical for G-protein binding, a few of these native amino-acids were introduced in the designed receptors. While these residues adopted only a partially active conformation in native agonist-bound A2AR structures^16^, all but Y288^7.53^ displayed a close to active conformation in the SPaDES_high7 structure (**Figure 4c-e**).

**Figure 4.**
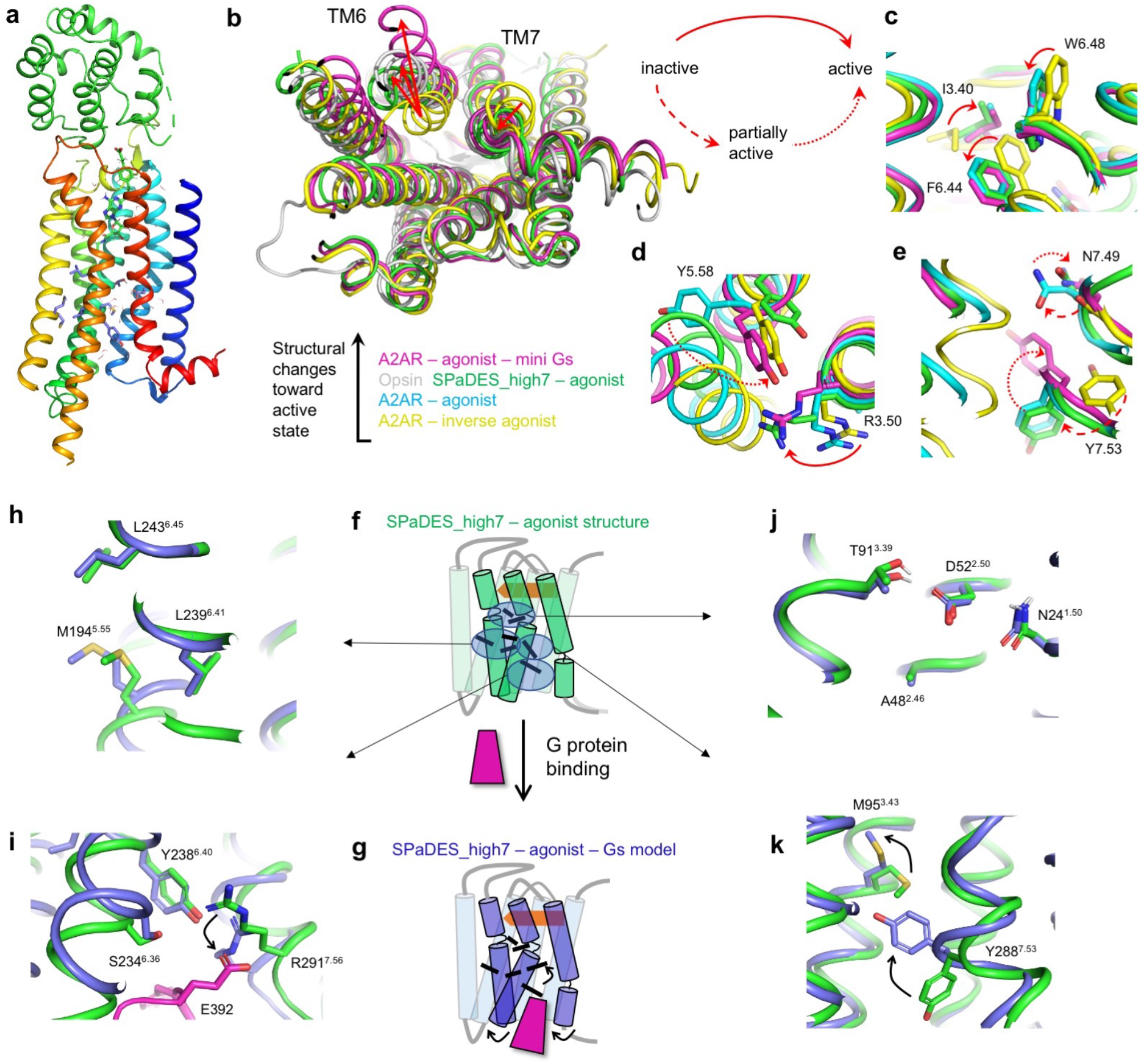
The designed SPaDES_high7 structure adopts an unforeseen activated conformation. **a.** Backbone representation of the designed SPaDES_high7 structure bound to the ligand agonist CGS21680 and N-terminally fused to a circularly permuted T4 lysozyme variant. **b.** Cytoplasmic view of the following superimposed structures: antagonist-bound A2AR (3eml, yellow), agonist-bound A2AR thermostabilized by alanine-scanning (2ydo, blue), agonist-bound computationally designed SPaDES_high7 (green), Gt peptide-bound opsin active state (3dqb, grey), agonist and mini-Gs bound A2AR thermostabilized by alanine-scanning (5g53, magenta). The structures are ranked based on the magnitude of conformational changes toward the fully active state (back arrow). **c-e.** Conformations of key conserved microswitches involved in receptor activation. **c.** TM3-5-6 interface around the W6.48 toggle switch in the receptor TM core. **d.** TM3-5-6 interface around the intracellular DR^3.50^Y motif. **e.** TM6-7 interface around the intracellular N^7.49^PxxY^7.53^ motif. Conformational changes between inactive, partially active and fully active states are indicated by distinct red arrows. **f-k.** Comparison between the conformation of the designed (black) and neighboring native (grey) residue microswitches in the agonist-bound designed SPaDES_high7 structure (**f**) and in the agonist and Gs-bound SPaDES_high7 design model (**g**). **h.** TM5-6 core interface. **i.** TM6-7 intracellular interface with bound Gs (magenta). **j.** TM1-2-3 core interface at the Na^+^ ion conserved binding site. **k.** TM3-6-7 intracellular interface.

The SPaDES_high7 structure was solved without bound Gs, while our active state design model included Gs was docked onto the G-protein binding site, generating specific receptor interactions and distortions on the intracellular sides of TM6 and TM7 (**Figure 4f-g**). Nevertheless, except for M95^3.43^ directly affected by the binding of Gs through the movements of the contacting Y288^7.53^ (**Figure 4k**), the conformations of the residues involved in the designed TMH interaction networks agreed with near-atomic accuracy between the predicted and experimental structures (**Figure 4h-i**). The RMSD between the experimental structure and the design model was only 1.5 Å over the entire TM region and reduced to 1.2 Å when two helical turns of TMH6 contacting Gs were excluded. Overall, the high accuracy structural prediction of the TMH core largely validates the *de novo* designed solvent-mediated allosteric interaction networks.

### *De novo* designed allosteric interaction networks define a new receptor active form

The SPaDES_high7 structure and function define a novel synthetic receptor active form that has not been observed in native GPCRs (**Figure 5**). We compared the designed features of SPaDES_high7 to the native properties of A2AR within the framework of a conformational energy landscape that integrates experimental evidence from X-ray, NMR spectroscopy and modeling calculations (**Figure 5a**). As revealed by seminal NMR experiments^17^, the WT A2AR adopts three main conformational states (inactive, partially, and fully active), with agonist binding stabilizing the partially active conformation. Several X-ray structures and functional studies of native GPCRs^4,18,19^ revealed the presence of a conserved binding site for a negative allosteric modulator Na^+^ ion molecule that stabilizes the inactive form. Na^+^ binding involves an intricate network of charged and neutral polar residues on TMHs 1, 2 and 3 (**Figure 5c,d**). Our calculations of ion occupancy suggest that the Na^+^ molecule binds to the inactive and partially active state conformations of A2AR, albeit with lower occupancy to the latter (**Figure 5a,e**). On the other hand, SPaDES_high7 mainly occupies an inactive, close to active (characterized by the above-described X-ray structure) and a fully active state conformation. In contrast to the WT receptor, the three states have similar energy in the apo form while agonist binding stabilizes the close to fully active state conformation. Our functional, structural data and calculations indicate only partial Na^+^ ion occupancy in the inactive state that becomes negligible in the two active states, where the volume of the designed polar cavity is too small to accommodate the ion molecule (**Figure 5b,d**).

**Figure 5.**
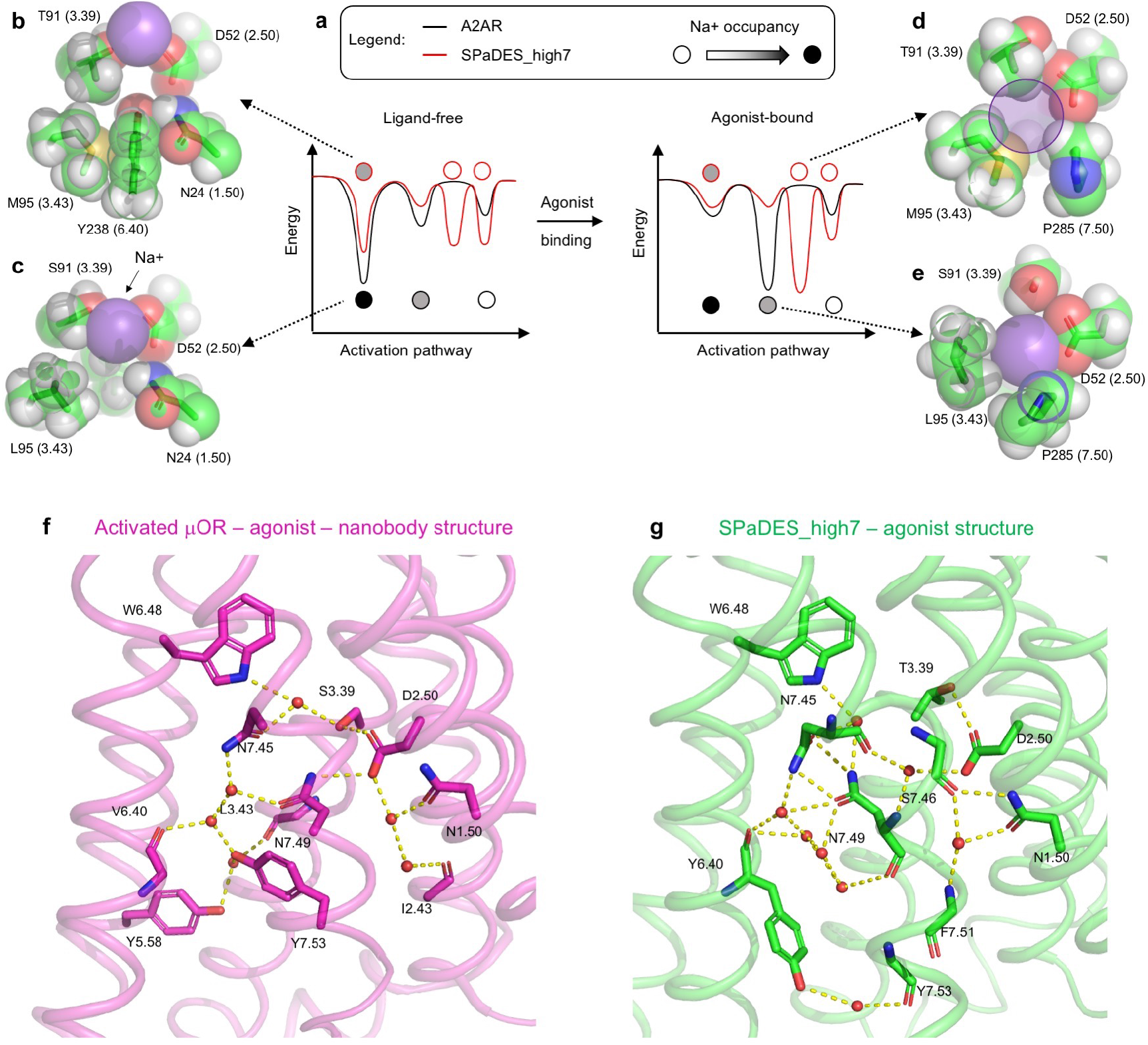
SPaDES-high7 is built of a synthetic allosteric interaction network distant from those of natural GPCRs. **a-e.** The designed core in SPaDES_high7 prevents the tight binding of the negative allosteric modulator Na^+^ ion and the receptor adopts a unique conformational energy landscape. **a.** Conformational energies calculated by SPaDES are reported for the A2AR (black) and SPaDES_high7 (red) receptors along the activation pathway without (left) and with bound agonist ligand (right). The transition energies were not calculated and their representation is fictive. Calculated Na^+^ ion occupancy is indicated using a filled circle and ranges from 0 (white) to 1.0 (black). **b-e.** Structure of the Na^+^ ion binding site in A2AR. The lowest energy position of the Na^+^ ion is indicated in purple sphere. Designed and WT amino-acids are in black and grey, respectively. **b.** Inactive state model of the designed SPaDES_high7. **c.** Inverse agonist ZM241385 bound A2AR X-ray (inactive state) structure (4eiy). **d.** Agonist CGS21680 bound SPaDES_high7 X-ray structure. The transparent purple sphere indicates the volume necessary for a Na^+^ ion to occupy the cavity. **e.** Partially active state adenosine bound A2AR X-ray structure (2ydo). **f-g.** Solvent-mediated polar interaction networks at static-switchable TMH interfaces in the μ-OR (5c1m) and SPaDES_high7 active state X-ray structures.

To directly assess how the designs rewired the solvent-mediated interaction networks defining the allosteric signal transduction pathways in receptor active states, we sought to compare the SPaDES_high7 structure with native receptor structures. Experimental evidence for solvent occupancy in GPCRs has been mostly derived from inactive state structures because of the fewer characterization and the often poor resolution of active state structures. However, the highest-resolution GPCR active state X-ray structure to date, i.e. that of mu-opioid (μOR) bound to agonist and nanobody, was solved at 2.1 Å resolution and enabled direct assignment of electron densities to buried solvent molecules^20^. For comparison with the μOR structure, the SPaDES_high7 structure was refined at atomic resolution using the software Phenix-SPaDES.refine that we developed to build and optimize protein models with explicit solvent against experimental crystallographic datasets. Phenix-SPaDES.refine considerably enhanced the quality of the SPaDES_high7 structural model as MolProbity, all-atom clash and *R*-free scores improved from 2.22, 3.72, 0.3086 to 1.19, 1.95 and 0.2978, respectively. All residues interacting with solvent molecules in the μOR structure are conserved in A2AR WT, thus justifying a direct comparison with the SPaDES_high7 structural model. The two structures revealed striking differences in the allosteric interhelical polar contact networks. A larger number of water molecules and hydrogen bond interactions were found in SPaDES_high7. However, the network in μOR bridged eight distinct TMH interfaces involving all TMHs except TMH 4. In contrast, polar connections in SPaDES_high7 mostly involved the designed static-switchable interfaces between TMHs 1, 2, 3 with TMHs 6 and 7, suggesting distinct allosteric mechanisms of receptor signaling (**Figure 5f,g**). In contrast to studies of natural GPCR sequences and structures^20,21^, our results reveal that structure stability and signaling activity can be designed concurrently through a wide range of interhelical polar contact networks, providing key allosteric coupling hotspots such as between the switchable TMH6 and TMH7 in GPCRs, efficiently propagate structural changes and movements.

## Discussion

Protein catalysis and allostery often rely on subtle protein motions and the complex interplay of protein-ligand, solvent cooperative interactions that remain very challenging to design. Here, we developed and applied a computational approach for engineering signaling activity into receptor scaffolds through the *de novo* design of solvent-mediated dynamic interaction networks at switchable TMH binding interfaces that are critical for signal transduction. Consistent with our predictions, we created receptors with a wide range of designed TMH interaction networks and measured signaling activities. Our most stable and active variant crystallized into an unforeseen active state conformation that matched the design model with near-atomic accuracy. Together with its high signaling activity and thermostability, SPaDES_high7 represents a new active form in the class of GPCRs.

Numerous structural and functional studies of native Rhodopsin-class GPCRs revealed a highly conserved network of allosteric polar residues buried in the TM region and interacting through water and ion molecules^3,20,22,23^. Alanine mutations of these sites often dramatically impaired signaling activities, demonstrating their critical involvement in signal transmission^19,24^. These results strongly suggested that efficient signaling functions rely on allosteric networks involving a precise and conserved topology of buried polar residues and water molecules. In contrast to these conclusions, we were able to engineer a library of signaling receptors through the *de novo* design of allosteric interaction networks that often bear limited similarity with those of naturally evolved receptors. The TMH and solvent interactions that mediate long-range communication pathways were considerably rewired in the designed receptors and defined novel network topologies (**Fig. 2, 5**). We also uncovered that the level of hydration at static switchable TMH interfaces is a strong unforeseen determinant of signaling efficacy and enabled the rational design of receptors with diverse responses. Together with the unique features of the SPaDES_high7 structure, these results highlight the high level of plasticity of “wet” polar interaction networks that critically promote conformational flexibility and allosterically propagate extracellular signals.

Our design calculations are only guided by the physics of protein interior solvation, the structural plasticity of microswitches and the stability of multiple protein conformational states. No pressure of selection for conservation of protein sequences is applied, and our *in silico* design approach is therefore agnostic of any evolutionary process that shape native protein sequences and structures. Our functional designs suggest that the space of protein interactions mediating protein movements and allostery is more extensive than what has been inferred from native proteins so far and remains largely unexplored. To our knowledge, our membrane receptors with synthetic cores provide the first evidence that protein allosteric functions can be rationally designed into protein scaffolds through *de novo* solvent-mediated interaction networks.

Protein-solvent and ligand molecule contacts represent a hallmark of protein catalysts and multi-pass membrane proteins, including transporters, channels and signaling receptors. Hence, by rationally designing solvent-mediated interaction networks and reprogramming long-range communication pathways, our approach paves the road for elucidating receptor signal transductions, catalytic mechanisms and engineering a wide diversity of novel protein functions impacting cell (e.g. cancer-immuno) therapies and synthetic biology. It should also prove particularly useful for accelerating the structure determination of challenging membrane protein targets by X-ray crystallography and cryoelectron microscopy, and the discovery of innovative drugs.

## Acknowledgements

The authors would like to thank members of the Barth lab for discussion, Joseph Ho, Charles Rauch, Marijane Russell, Michael Sauder, Sheela Ashok, Aiping Zhang, and Cheyenne Logan from the Lilly Structural Biology group for technical support and helpful discussions, and Mark Bures for helping initiate the project. K-Y.C. was partially funded by a training fellowship from NIH NIGMS T32GM008280. This work was supported by Eli Lilly and Company through the Lilly Research Award Program (LRAP), a grant from the National Institute of Health (1R01GM097207), a grant from the Swiss National Science Foundation (31003A_182263), and a supercomputer allocation from XSEDE (MCB120101) to P.B. P.B. was also supported by funding from EPFL and the Ludwig Institute for Cancer Research. This research used resources of the Advanced Photon Source, a U.S. Department of Energy (DOE) Office of Science User Facility operated for the DOE Office of Science by Argonne National Laboratory under Contract No. DE-AC02-06CH11357. Use of the Lilly Research Laboratories Collaborative Access Team (LRL-CAT) beamline at Sector 31 of the Advanced Photon Source was provided by Eli Lilly Company, which operates the facility. We thank Diamond Light Source for beamtime (Proposal #IN14323-4) and the staff of beamlines I04-1 for assistance with crystal testing and data collection.

## Author Contributions

P.B., H.E., K.B. designed the studies. J.L. developed the computational methods. K-Y.C. and J.L. performed the calculations. K-Y.C. performed functional experiments on SPaDES designs. A.R. designed the T4Lcp-SPaDES_high chimeric constructs. X.Z., B.C. performed SPaDES_high protein purification. K.B., J.W., F.T., L.K., assayed thermostability, conducted crystallization studies, and solved the SPaDES_high7 X-ray structure. J.B. processed X-ray crystallization data. All authors analyzed the data. P.B. wrote the manuscript.

## Methods

### SPaDES gene construction and expression

HA-tagged A2AR gene in pcDNA3.1(+) was obtained from the cDNA library. Genes coding for SPaDES variants were synthesized. Plasmids were transiently transfected using Genejuice (Novagen/EMD Millipore) into HEK293T cells. HEK293T cells at 70-80% confluency in 15cm tissue culture plates were transfected with 4ug of DNA per plate. After 24 hours, transfection medium was replaced with standard growth medium (DMEM supplemented with L-glutamine (2mM), penicillin (100μg/mL), streptomycin (100μg/mL), and fetal bovine serum (10%) and cells were grown for an additional 24 hours prior to harvesting.

### Membrane preparation

Membranes were prepared from transfected cells using sucrose gradient centrifugation as previously described. Briefly, cells from 10cm plates were collected by cell scraper with PBS solution. Cells were pelleted and resuspended in cold hypotonic buffer (1mM Tris-HCL, pH 6.8, 10mM EDTA, protease inhibitor cocktail). Cells were forced through a 26-gauge needle three times. The cell lysate was layered onto a 38% sucrose solution in buffer A (150mM NaCl, 1mM MgCl, 10mM EDTA, 20mM Tris-HCl, pH 6.8, protease inhibitor cocktail) in SW-28 ultracentrifuge tubes. Cells were centrifuged at 15,000 rpm at 4°C for 20 minutes, followed by collection of the interface band with an 18-gauge needle. The collected solution was transferred to Ti-45 ultracentrifuge tubes (Beckman) and the volume brought up to 50mL with buffer A. The sample was spun at 40,000 rpm at 4°C for 30 minutes. The membrane pellets from each 10cm plate was resuspended in 0.5mL buffer A and stored at −80°C in 100μL aliquots.

### SPaDES designs partial purification

SPaDES variants were partially purified from thawed membrane preparations immediately prior to assaying via anti-HA agarose beads. Membrane preparations were solubilized with 1% n-dodecylmaltoside (DDM) for one hour at 4°C, and loaded onto anti-HA agarose beads for one hour at 4°C. The beads were washed with TBS with 0.1% DDM wash buffer three times and HA-tagged receptor variants were eluted with HA peptide (1mg/mL in TBS with 0.1% DDM). Purity of affinity purified SPaDES samples was assessed by coomassie-stained SDS-PAGE along with western blots for all receptor variants.

### Expression and purification of the G protein Gs

G-alphaS (Gas) was cloned into pFastbacI followed by transformation into DH10α cells. Recombinant bacmid DNA was isolated and transfected in Sf9 insect cells with Cellfectin II. The transfected cells were grown at 28°C for 72 hours followed by centrifugation in 15mL Falcon tubes to pellet cell debris. The supernatant was saved as P1 viral stock. Virus was amplified by infecting Sf9 cells with P1 viral stock solution at a 2-fold multiplicity-of-infection and cultured for 72 hours prior to harvesting. This amplification process was repeated to generate a high-titer P3, which was used for infection and protein expression. P3 stock was used to infect Sf9 cells at an MOI greater than 4 and harvested 48-60 hours post infection. Cells were washed three times in ice-cold PBS and resuspended in homogenization buffer (10mM Tris-HCl, 25mM NaCl, 10mM MgCl_2_, 1mM EGTA, 1mM DTT, protease inhibitor cocktail, 10μM GDP, pH 8.0). GaS was purified as described previously^25^. Antibodies were purchased from Santa Cruz Biotechnology.

### G protein activation assays

Receptor variants were assayed for their ability to induce guanine nucleotide exchange by the GaS. To measure constitutive activity the reaction mixture consisted of 4μM GaS, 20uM of [^35^S]-GTPgS mix, 50mM Tris-HCl, pH 7.2, 100mM NaCl, 4mM MgCl_2_, 1mM dithiothreitol. Receptor concentrations were ~5-20nM per sample as estimated from absorbance at 280nm of the anti-HA agarose purified samples. For all samples used for reactions, western blots were performed to normalize for receptor quantity using monoclonal anti-HA antibody (Thermo Scientific). Reactions were started by adding 150μL partially purified receptor samples to 300μL of reaction mixture and incubating on ice for 1 hour. To measure ligand induced receptor activities a saturating concentration of adenosine (10μM) was added to the reaction mixture. Reactions were stopped by filter binding onto Millipore nitrocellulose filters. Filters were washed three times with ice-cold TBS prior to incubation with scintillation fluid. Radioactivity counts were measured on a Beckman LS6000 scintillation counter. Mock transfected samples (using empty pcDNA vector) and WT receptor served as controls. Background binding measured by mock transfected samples was subtracted and activities of receptor variants were normalized relative to WT receptor via densitometry analysis of western blots using ImageJ software. Statistical significance of differences in constitutive or ligand-induced activities was assessed by student t-tests using GraphPad software.

### Apparent agonist-bound SPaDES stability

Apparent stability of agonist-bound SPaDES variants was determined by measuring either receptor binding to agonist (apparent melting temperature) or receptor activities (apparent receptor half-life) as a function of temperature. To measure apparent melting temperature, anti-HA agarose affinity purified receptor samples prepared as described above were first pre-incubated at increasing temperatures for 30 min. The samples were then incubated with saturating amount of [^3^H]-adenosine (10μM) in a total volume of 150μL per sample for 1 hour on ice followed by loading onto Millipore nitrocellulose washed for scintillation counting as mentioned above. To measure apparent receptor half-life, purified receptor samples were pre-incubated at 37 °C over a time period from 0-60 minutes prior to addition to the reaction mixture containing GaS, [^35^S]-GTPγS and 10μM adenosine as described above in measuring G protein activation. Apparent melting temperatures and active state half-life curves were fitted using GraphPad Prism software and analyzed for statistical significance by student t-tests.

### Modeling of SPaDES active state conformation

The modeling was performed before the release of the A2AR active state structure bound to mini-Gs^13^. Hence we modeled SPaDES active conformation by homology to the fully active Gs-bound beta 2 adrenergic receptor (B2AR) structure^26^. Since the sequences of B2AR and A2AR on the extracellular side diverge substantially, we took advantage of available partially active agonist-bound A2AR structures. We grafted the extracellular region of UK-432097 agonist-bound A2AR structure^27^ onto the intracellular region of the fully active homolog Gs-bound (B2AR) X-ray structure to create a chimeric homolog template for the fully active conformation of the agonist and Gs-bound SPaDES template. Intracellular loop regions were rebuilt *de novo* in the presence of bound Gs as described in^28^, and the entire structure was relaxed using SPaDES, a scoring function developed for modeling membrane proteins with explicit solvent^10^. Around 20000 independent simulations were performed and, from those, the 10% lowest energy models were clustered, with thecenters of the most populated clusters selected as fully active state SPaDES templates for the design calculations of activating microswitches. The A2AR inactive state X-ray structure bound to the high-affinity antagonist ZM241385^29^ was selected as a representative inactive state model for the SPaDES design templates.

### Expression vector design of the T4L fusion construct

There are numerous, successful examples of T4 lysozyme (T4L) functioning as a stabilizing chaperon protein for numerous GPCR proteins deposited in the pdb database. Our strategy was to follow those successful examples and fuse a cysteine-free T4L^30^ to the N-terminal of A2AR with one exception; we decided to use a circular permutant of T4L (T4Lcp) designed by Cellitti *et al*. that relocates the N-terminal A-helix to the C-terminal, creating subdomains that are contiguous in sequence and are linked by G4S sequence^31^. T4Lcp crystallized at higher resolution (1.8 Å) than other T4L constructs and may therefore have a higher probability to positively impact SPaDES crystallization. The T4Lcp – SPaDES fusion sequence was optimized using the structure of the circular permutant T4L protein (pdb code 2o7a) as follows. To link the two proteins, we designed small linkers between the C-terminus of T4Lcp and the N-terminal amino acid of SPaDES_high variants. Following *in silico* screening and experimental testing, the two amino acid linker Ala-Pro was found to be the most successful construct that resulted in a stable, soluble protein and foremost, the one that crystallized.

### Cloning and Baculovirus expression of T4Lcp-SPaDES_high variants

A circular permutant T4 lysozyme 207A (T4Lcp)-SPaDES_high7 fusion including A2AR encompassing residues 2-316 with designed mutations L48A, S91T, I238Y, L194M, V239L, A243L, L95M (numbered relative to reference sequence NP_000666) was PCR amplified and TOPO cloned into a custom TOPO adapted pFastBac (KX) vector (Life Technologies). cDNA from this clone was named 14022b11KXem1h1 and sequence verified. Expression of this vector generated MKTIIALSYI FCLVFADYKD DDDGAP/ T4L 207A/ AP/A2AR aa2-316 plus L48A, S91T, I238Y, L194M, V239L, A243L, L95M /GS/LVPRGS/HHHHHHHHHH. T4Lcp 207A corresponds to amino acids 2-124 (numbered relative to reference sequence NP_049736). Standard baculovirus expression using a modified version of the Bac to Bac system protocol (Life Technologies) in combination with the DH10EMBacY bacmid (Geneva Boiotech) was used to generate the virus. T4Lcp-A2AR constructs were expressed in S*f*9 cells which were cultured for 48 hours, harvested by centrifugation and pelleted for storage at −80°C prior to purification.

### T4Lcp-SPaDES_high7 Purification

Protein purifications were carried out using cobalt affinity, thrombin cleavage, PNGase F deglycosylation and size-exclusion chromatography (SEC). Infected Sf9 cell pellets were lysed in 20 mM Tris, pH 7.5, 500mM NaCl, 0.2mM MNGC-12, and 1mM CGS21680. Membrane samples were collected by spinning at 44,000 × rpm for 45min. Extraction of T4Lcp-SPaDES_high7 from Sf9 membranes was done with a Dounce homogenizer in a solubilization buffer comprised of 20mM *n*-dodecyl β-d-maltoside (DDM), 4mM cholesteryl hemisuccinate (CHS), 50 mM Hepes, pH 7.8, 500 mM NaCl, 25 μM of Cmpd-1. Talon resin was added and mixed continuously for overnight at 4 °C. The Talon resin was collected by spinning, and washed extensively with Ni wash buffer comprised of 5mM DDM, 1mM CHS, 50 mM Hepes, pH 7.8, 500 mM NaCl, 10 mM imidazole. This was followed by a second washing step using 2mM MNGC-12 instead of DDM, CHS; 50 mM Hepes, pH 7.8, 500 mM NaCl, 10 mM imidazole. The protein was eluted from the resin using the elution buffer comprising 250 mM imidazole, 1mM MNGC-12, 50 mM Hepes, pH 7.8, 500 mM NaCl. A desalting column was then used to exchange buffer into 50mM Hepes pH 7.8, 500mM NaCl,0.5mM MNGC-12, 0.1mM CGS21680, following by 20μl thrombin and 20μl PNGase F for overnight 4 °C cleavage. The elution sample was concentrated and loaded onto a SEC column (Superdex 200) with the buffer 0.2mM MNGC-12, 20 mM Hepes, pH 7.5, 500 mM NaCl, and 25 μM CGS21680. The major peak fractions were combined and concentrated for crystallization and the ligand was added up to 500 μM final concentration.

### Fluorescent Thermal Stability Assay of T4Lcp-SPaDES_high7 constructs

Receptors were purified as described for crystallization with reducing agent removed in the final chromatography step. Receptors were diluted to 2μM into PBS, 0.5mM DDM, 0.1mM CHS, 20μM (7-Diethylamino-3-(4’-Maleimidylphenyl)-4-Methylcoumarin), 100μM ligand compound. Samples were incubated on ice for 1 hour prior to measurement. Melts were completed in a Qiagen Rotor-Gene Q monitoring on the blue channel with a thermal ramp from 30-90 °C measuring fluorescence 3 times per degree. Data was processed using Excel. Raw fluorescence was normalized from which melting temperatures (Tms) were calculated.

### T4Lcp-SPaDES_high7 crystallization

T4Lcp-SPaDES_high7 protein (50mg/ml) with 0.5mM of the ligand agonist CGS21680 complex was mixed with monoolein containing 10% cholesterol in 1:1.5 parts v/v protein:lipid ratio using the twin-syringe mixing method^32^. We performed extensive LCP crystallization trials for T4Lcp-SPaDES_high7 with a mosquito^®^ robot (TTP Labtech) at Room Temperature. Initial crystals came from conditions of 0.1M Hepes pH 7-8, 25-30% PEG300, 50-200mM sodium potassium tartrate, two days after LCP trays were set up. Optimization screens were performed with well buffer (0.1M Hepes pH 7, 27% PEG300, 50mM sodium potassium tartrate) with 10% detergent screen (Hampton Research). Final crystals datasets were collected to 3.9Å resolution using conditions of 0.1M Hepes pH 7, 27% PEG300, 50mM sodium potassium tartrate with 5mM CYMAL-5.

### Data collection, processing and structure determination

A complete X-ray data set was collected on a single rod-shaped crystal (~ 10 × 100 μm2) at 100K in a single sweep of 900 × 0.2° oscillations and 0.4 s exposure at DLS-I04-1 at Diamond Light Source in Harwell, UK. The whole sample (i.e. the LCP sample/blob in the MiTeGen loop that contained the crystal) was exposed to the beam during data collection. The diffraction data were integrated using autoPROC/XDS^33,34^ and merged and scaled in SCALA^35^ from the CCP4 suite^36^. The crystal structure of the T4Lcp-SPaDES_high7-CGS21680 complex was determined by molecular replacement using the previously solved T4Lcp and A2AR structures: 2o79^31^ and modified 5IU4^37^ as separate search models. Clear density for the ligand was observed immediately. After numerous cycles of refinement with REFMAC5^38^, the model was further refined using Phenix-SPaDES.refine as described below to reach reasonable *R* factors and MolProbity scores.

## References

1. Changeux, J.P. & Christopoulos, A. Allosteric Modulation as a Unifying Mechanism for Receptor Function and Regulation. Cell 166, 1084–1102 (2016).

2. Yuan, S., Hu, Z., Filipek, S. & Vogel, H. W246(6.48) opens a gate for a continuous intrinsic water pathway during activation of the adenosine A2A receptor. Angew Chem Int Ed Engl 54, 556–9 (2015).

3. Yuan, S., Filipek, S., Palczewski, K. & Vogel, H. Activation of G-protein-coupled receptors correlates with the formation of a continuous internal water pathway. Nat Commun 5, 4733 (2014).

4. Ye, L. et al. Mechanistic insights into allosteric regulation of the A2A adenosine G protein-coupled receptor by physiological cations. Nat Commun 9, 1372 (2018).

5. Setny P., W.M.D. Water-mediated conformational preselection mechanism in substrate binding cooperativity to protein kinase A. PNAS 115, 3852–3857 (2018).

6. Lu, P. et al. Accurate computational design of multipass transmembrane proteins. Science 359, 1042–1046 (2018).

7. Scott E Boyken 1, Z.C., Benjamin Groves 3, Robert A Langan 4, Gustav Oberdorfer 5, Alex Ford 4, Jason M Gilmore 4, Chunfu Xu 4, Frank DiMaio 4, Jose Henrique Pereira 6, Banumathi Sankaran 7, Georg Seelig 8, Peter H Zwart 9, David Baker 10 De novo design of protein homo-oligomers with modular hydrogen-bond network-mediated specificity. Science 352, 680–7 (2016).

8. T M Jacobs 1, B.W., T Williams 2, X Xu 3, A Eletsky 3, J F Federizon 4, T Szyperski 4, B Kuhlman 5. Design of structurally distinct proteins using strategies inspired by evolution. Science 352, 687–90 (2016).

9. Jiayi Dou # 1 2, A.A.V., William Sheffler 1 2, Lindsey A Doyle 3, Hahnbeom Park 1 2, Matthew J Bick 1 2, Binchen Mao 1 4, Glenna W Foight 5, Min Yen Lee 5, Lauren A Gagnon 5, Lauren Carter 1 2, Banumathi Sankaran 6, Sergey Ovchinnikov 1 2 7, Enrique Marcos 1 2 8, Po-Ssu Huang 1 2 9, Joshua C Vaughan 5, Barry L Stoddard 3, David Baker 10 11 12. De novo design of a fluorescence-activating β-barrel. Nature 561, 485–491 (2018).

10. Lai, J.K., Ambia, J., Wang, Y. & Barth, P. Enhancing Structure Prediction and Design of Soluble and Membrane Proteins with Explicit Solvent-Protein Interactions. Structure 25, 1758–1770 e8 (2017).

11. Magnani, F., Shibata, Y., Serrano-Vega, M.J. & Tate, C.G. Co-evolving stability and conformational homogeneity of the human adenosine A2a receptor. Proc Natl Acad Sci U S A 105, 10744–9 (2008).

12. Scheerer, P. et al. Crystal structure of opsin in its G-protein-interacting conformation. Nature 455, 497–502 (2008).

13. Carpenter, B., Nehme, R., Warne, T., Leslie, A.G. & Tate, C.G. Structure of the adenosine A(2A) receptor bound to an engineered G protein. Nature 536, 104–7 (2016).

14. Ahuja, S. & Smith, S.O. Multiple switches in G protein-coupled receptor activation. Trends Pharmacol Sci 30, 494–502 (2009).

15. Nygaard, R., Frimurer, T.M., Holst, B., Rosenkilde, M.M. & Schwartz, T.W. Ligand binding and micro-switches in 7TM receptor structures. Trends Pharmacol Sci 30, 249–59 (2009).

16. Lebon, G. et al. Agonist-bound adenosine A2A receptor structures reveal common features of GPCR activation. Nature 474, 521–5 (2011).

17. Ye, L., Van Eps, N., Zimmer, M., Ernst, O.P. & Prosser, R.S. Activation of the A2A adenosine G-protein-coupled receptor by conformational selection. Nature 533, 265–8 (2016).

18. Liu, W. et al. Structural basis for allosteric regulation of GPCRs by sodium ions. Science 337, 232–6 (2012).

19. Massink, A. et al. Sodium ion binding pocket mutations and adenosine A2A receptor function. Mol Pharmacol 87, 305–13 (2015).

20. Huang, W. et al. Structural insights into micro-opioid receptor activation. Nature 524, 315–21 (2015).

21. Venkatakrishnan, A.J. et al. Molecular signatures of G-protein-coupled receptors. Nature 494, 185–94 (2013).

22. White, K.L. et al. Structural Connection between Activation Microswitch and Allosteric Sodium Site in GPCR Signaling. Structure 26, 259–269 e5 (2018).

23. Angel, T.E., Chance, M.R. & Palczewski, K. Conserved waters mediate structural and functional activation of family A (rhodopsin-like) G protein-coupled receptors. Proc Natl Acad Sci U S A 106, 8555–60 (2009).

24. Nygaard, R., Valentin-Hansen, L., Mokrosinski, J., Frimurer, T.M. & Schwartz, T.W. Conserved Water-mediated Hydrogen Bond Networkbetween TM-I, −II, −VI, and-VII in 7TM Receptor Activation. Journal of Biological Chemistry 285, 19625–19636 (2010).

25. Graber, S.G., Figler, R.A. & Garrison, J.C. Expression and purification of functional G protein alpha subunits using a baculovirus expression system. J Biol Chem 267, 1271–8 (1992).

26. Rasmussen, S.G. et al. Crystal structure of the beta2 adrenergic receptor-Gs protein complex. Nature 477, 549–55 (2011).

27. Xu, F. et al. Structure of an agonist-bound human A2A adenosine receptor. Science 332, 322–7 (2011).

28. Chen, K.Y., Sun, J., Salvo, J.S., Baker, D. & Barth, P. High-resolution modeling of transmembrane helical protein structures from distant homologues. PLoS Comput Biol 10, e1003636 (2014).

29. Jaakola, V.P. et al. The 2.6 angstrom crystal structure of a human A2A adenosine receptor bound to an antagonist. Science 322, 1211–7 (2008).

30. Matsumura, M. & Matthews, B.W. Control of enzyme activity by an engineered disulfide bond. Science 243, 792–4 (1989).

31. Cellitti, J. et al. Exploring subdomain cooperativity in T4 lysozyme I: structural and energetic studies of a circular permutant and protein fragment. Protein Sci 16, 842–51 (2007).

32. Caffrey, M. & Cherezov, V. Crystallizing membrane proteins using lipidic mesophases. Nat Protoc 4, 706–31 (2009).

33. Vonrhein, C. et al. Data processing and analysis with the autoPROC toolbox. Acta Crystallogr D Biol Crystallogr 67, 293–302 (2011).

34. Kabsch, W. Integration, scaling, space-group assignment and post-refinement. Acta Crystallogr D Biol Crystallogr 66, 133–44 (2010).

35. Evans, P. Scaling and assessment of data quality. Acta Crystallogr D Biol Crystallogr 62, 72–82 (2006).

36. Winn, M.D. et al. Overview of the CCP4 suite and current developments. Acta Crystallogr D Biol Crystallogr 67, 235–42 (2011).

37. Segala, E. et al. Controlling the Dissociation of Ligands from the Adenosine A2A Receptor through Modulation of Salt Bridge Strength. J Med Chem 59, 6470–9 (2016).

38. Murshudov, G.N. et al. REFMAC5 for the refinement of macromolecular crystal structures. Acta Crystallogr D Biol Crystallogr 67, 355–67 (2011).

